# A chemically defined oviposition attractant and repellent of Black Soldier Flies (*Hermetia illucens*)

**DOI:** 10.1101/2024.06.04.597456

**Authors:** Nyasha KT Thomas, Zsolt Karpati, Thomas Schmitt, Olena Riabinina

## Abstract

Black Soldier Flies (BSF), *Hermetia illucens*, are industrially important species. They can consume large amounts of spoilt organic material as larvae and bio-convert it to more useful biomass. Female BSF lay eggs in crevices adjacent to spoilt organic materials that serve as an oviposition attractant. These kairomones are central to maximising rearing efforts, yet the composition and origin of oviposition cues remain undefined, and no synthetic oviposition attractants are currently available. This work aimed to identify key components of naturally occurring oviposition attractants and to formulate an effective synthetic alternative for BSF. We have developed a novel oviposition assay and found larval food- and frass-based attractants to be the most effective at centralizing egg laying. We have identified the volatile compounds in the headspaces of putative attractants and established that the antennae of the female flies respond to some of these compounds. Behavioural validation using synthetic compounds allowed us to generate a mixture of 5 compounds (p-cresol, decanal, sulcatone, pentanoic acid, acetophenone) that cues oviposition as efficiently as currently used natural oviposition attractants. We also identified a synthetic mixture that deters oviposition in BSF. The synthetic attractant and repellent we generated are likely to simplify BSF rearing in research and industrial settings.

## Introduction

The rising demand for high-protein animal feeds in agriculture and the increased production of organic waste in the industry are two problems that can be minimized by one organism. Black Soldier Flies (BSF), *Hermetia illucens* L. (Diptera: Stratiomyidae), originate from South America, but can now be found internationally due to their usefulness in farming (e.g. Kaya et al., 2021). During their larval stage, they consume spoilt organic materials and essentially bio-convert them into useful biomass. Dried larvae of this species are one of only seven insect species approved by the European Commission to be used as compound feed for aquaculture, with more expansive legislation in Canada and the US utilizing them in pet foods (Diener et al., 2011; Ferrarezi et al., 2016). BSF are polyphagous, able to consume a large variety of organic materials from common food scraps to putrefied *S. grandiflora* leaves (Kuppusamy et al., 2020; Tschirner and Simon, 2015). Larvae possess high nutritional value, particularly regarding their protein and fat content, reaching over 42% and 37% respectively, therefore are primarily exploited as animal feed (Chia et al., 2018; Spranghers et al., 2017). Additionally, their fats can be used as raw material for biodiesel and bioplastic production, as well as their frass waste can be utilized as fertilizers (Barbi et al., 2021; Zheng et al., 2013). Despite the growing popularity of this species within large industries, and thus the demand for mass rearing, the behaviours and biology of BSF are yet to be properly characterised.

Successful mating of BSF adults occurs between females two days post-emergence or older and males who eclose sexually mature (Julita et al., 2020). If mating is successful and fertilization occurs, gravid females will lay clutches of eggs in small crevices adjacent to decomposing organic materials (Hoc et al., 2019). Moisture, substrate composition and temperature affect BSF oviposition (Julita et al., 2020; Tomberlin and Sheppard, 2002). Increased temperatures, humidity and nutritionally dense feed have all been shown to accelerate larval growth and adult lifespan and have been suggested to marginally influence mating, rates of ovipositing and egg clutch size (Julita et al., 2020; Sripontan et al., 2017; Tomberlin and Sheppard, 2002).

BSF may be encouraged to deposit their eggs at one specific location by using volatile oviposition attractants. Specific organic materials may function as attractants, wherein gravid females will lay eggs in adjacent crevices with varying intensity (Park et al., 2016; Tomberlin and Sheppard, 2002). Specific natural attractants with complex bouquets, such as manure and fruit waste, have been identified, however, their attractive compounds remain unknown (Chin Tan and Sim, 2020; Nyakeri et al., 2017). For example, Nyakeri *et al*. (2017) found that cow manure was the most consistent stimulant, attracting the highest weight of laid eggs. They also more generally found that rotting materials such as frass were more effective at attracting appreciable egg laying than unspoilt materials, hypothesising that BSF females are attracted to volatiles associated with decomposition. Other studies have led to contrasting results, some reporting fruit waste to be a better attractant than other food wastes or manure, indicating that different strains of BSF may be attracted to different oviposition substrates (Sripontan et al., 2017). Conspecific eggs are also thought to act as oviposition cues via the emission of volatiles (Zheng et al., 2013). The authors of this study further investigated the specific cues by comparing the attractiveness of egg clutches with sterile eggs and no eggs, finding that egg sterilization resulted in decreased deposition and that lack of eggs was the least attractive (Zheng et al., 2013). Additionally, a higher response was recorded when a mixture of multiple bacteria species was present compared to single isolated strains. This suggests that ovipositing cues from conspecific eggs have a microbial origin. On the other hand, the increasing presence of bacteria originating from competitors of the BSF, such as the lesser mealworm, led to decreased ovipositing (Zheng et al., 2013). This points to olfactory cues as not only indicating potential food sources for BSF offspring but also for avoiding competitor species. However, the specific volatiles that induce BSF to lay eggs are currently unknown, and synthetic oviposition attractants are not available.

In this study, we identified key volatiles that are present in natural oviposition attractants of BSF. To test the efficacy of our newly identified compounds, we formulated a synthetic oviposition attractant and an oviposition repellent. The availability of the synthetic attractant and repellent may prove useful for laboratory and industrial rearing of BSF, as they are easy to prepare and do not require handling of decomposing organic materials.

## Materials and Methods

### Colony maintenance

*Hermetia illucens* were obtained from BetaBugs Ltd., Edinburgh, UK, and maintained in a climate-controlled room at 27 °C and 70-75% relative humidity (RH), on an 8:16 hour light:dark cycle. The feed for the larvae consisted of a 10:1 mixture of chick crumb (Mr Johnson’s Free Range Chick Crumb, Mr Johnson’s Petcare Ltd., UK): white sugar that was supplemented with water to reach a semi-liquid texture. The 1^st^ instar larvae and the feed were placed in 210 ml tamper-proof containers (#4506, Jars & Things, UK). The 210 ml containers were used for initial larval growth to minimise mould growth on the food substrate. Holes were cut in the lids of the tamper-proof containers and the holes were covered with 30 gsm garden fleece (ASIN:B073YTD83Y, Jagmaronline) to allow for air exchange. Once larvae reached 3-4^th^ instar, they were transferred to larger 1140 ml tamper-proof containers (#4523, Jars & Things, UK) with additional fresh food and covered with cut-out lids with garden fleece. As containment measures, larval containers were placed in large plastic buckets filled with water to 4-6 cm depth to prevent larval escape, and garden fleece in the lid was changed when it appeared soiled.

After 15-21 days, larvae darkened in colour as pupation began. Once larvae began to pupate, they were cleaned with water from any remaining food, dried, placed in clean 870 ml tamper-proof containers with fleece cut-outs (#4516, Jars & Things, UK) and left to pupate. Once larvae reached the pupal stage, the tamper-proof containers were placed without lids in fabric cages (32.5 cm x 32.5 cm x 32.5cm, BugDorm, MegaView Science Co., Ltd., Taiwan) and the pupae were left for the imagines to emerge within the cages. The pot of pupae was moved to new clean empty cages every day, ensuring that each cage was inhabited by adults that eclosed on the same day. Even-aged adults were kept in cages and sprayed with water daily, with no food provided. Adult flies lived between 8 and 15 days.

### Egg collection

Adult female BSF became sexually mature 2-4 days after emergence and were observed mating only after this time, whereas males were sexually mature when they emerged (Giunti et al., 2018; Julita et al., 2020; Tomberlin et al., 2002). Gravid BSF females began laying eggs 2 days after mating, which corresponds to 4 to 10 days post-emergence (Julita et al., 2020; Liu et al., 2020). Acrylic egg traps (provided by BetaBugs) that were designed to mimic crevices in wood were placed atop the prepared attractant pots, with gravid females searching for these artificial crevices using their extended ovipositors to lay eggs. Egg traps were removed daily and replaced with fresh traps, eggs within the egg traps were removed, weighed, and stored in 5 ml plastic pots for 1-3 days until the larvae hatched.

### Sex determination

Adult BSF were sexed using the anatomical dimorphism at the endpoint of the abdominal segments. Females possess sharp, forked abdominal tips, functioning as both genital organs and oviducts. The abdominal endpoints of males have a hook-like structure as well as an aedeagus (Malawey et al., 2019).

### Natural oviposition attractants

Five different substrates were tested as oviposition attractants in behavioural assays (***Table 1***). Attractants were made-up at a 1:10 substrate:deionised water ratio, left for 24 hours at 27 °C and 70-75 % RH, then frozen in aliquots until use. 25 ml of attractant and 25 ml of deionised water control were placed in two separate 210 ml tamper-proof tubs, with mesh cut-outs in the lids that allowed attractant volatiles to emanate, but blocked flies from coming into physical contact with the attractant substrates. Egg traps were placed atop attractant pots during egg collection and behavioural assays.

**Table 1.** Putative attractants and their corresponding solvent controls.

| Attractant | Composition and Description | Solvent Control |
| --- | --- | --- |
| Eggs | Egg clutches collected from conspecifics | n/a |
| BDB | BSF larval and pupal remains brewed in water | Deionised Water |
| Dry | Remaining substrate and waste from larvae fed on chick crumb and spent brewing grains, rehydrated. Provided by BetaBugs Ltd. | Deionised Water |
| Frass | Remaining substrate and waste from larvae fed on chick crumb, rehydrated | Deionised Water |
| Food | Larval feed substrate consisting of a 10:1 ratio of chick crumb and white sugar, hydrated | Deionised Water |

### Binary-choice behavioural assays

Even aged females (median number of females/assay across all assays: 43, 1^st^ quartile: 24, 3^rd^ quartile: 74) and males, paired in approx. 1:1 ratio, were observed until mating occurred (2-4 days after emergence). Once mating had been observed, adults were entered into the behavioural assay. A T-maze constructed from cylindrical plastic pots connected a starting cage containing mated adults to two cages each containing an attractant pot and egg trap (***Figure 1***). All three cages were 17.5 cm x 17.5 cm x 17.5 cm (BugDorm, MegaView Science Co., Ltd., Taiwan). Flies were able to move freely between the three cages via the plastic tunnel during the duration of the closed 4-day assay. The eggs were collected and weighed daily, and the new egg trap was provided daily. The same attractant pots and egg traps were used throughout the assay. To evaluate the success of the assay, the total eggs per female ratio was calculated 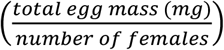, and assays were considered successful if the egg mass per female exceeded 2 mg. If the mass was less than this, it was assumed that the mating was unsuccessful, and the trial was discarded. For all trials, adults were euthanised via freezing (-20 °C) after the assay, then counted and sexed, whilst eggs were collected daily and weighed, then either fed upon hatching as described above or frozen and discarded.

**Figure 1.**
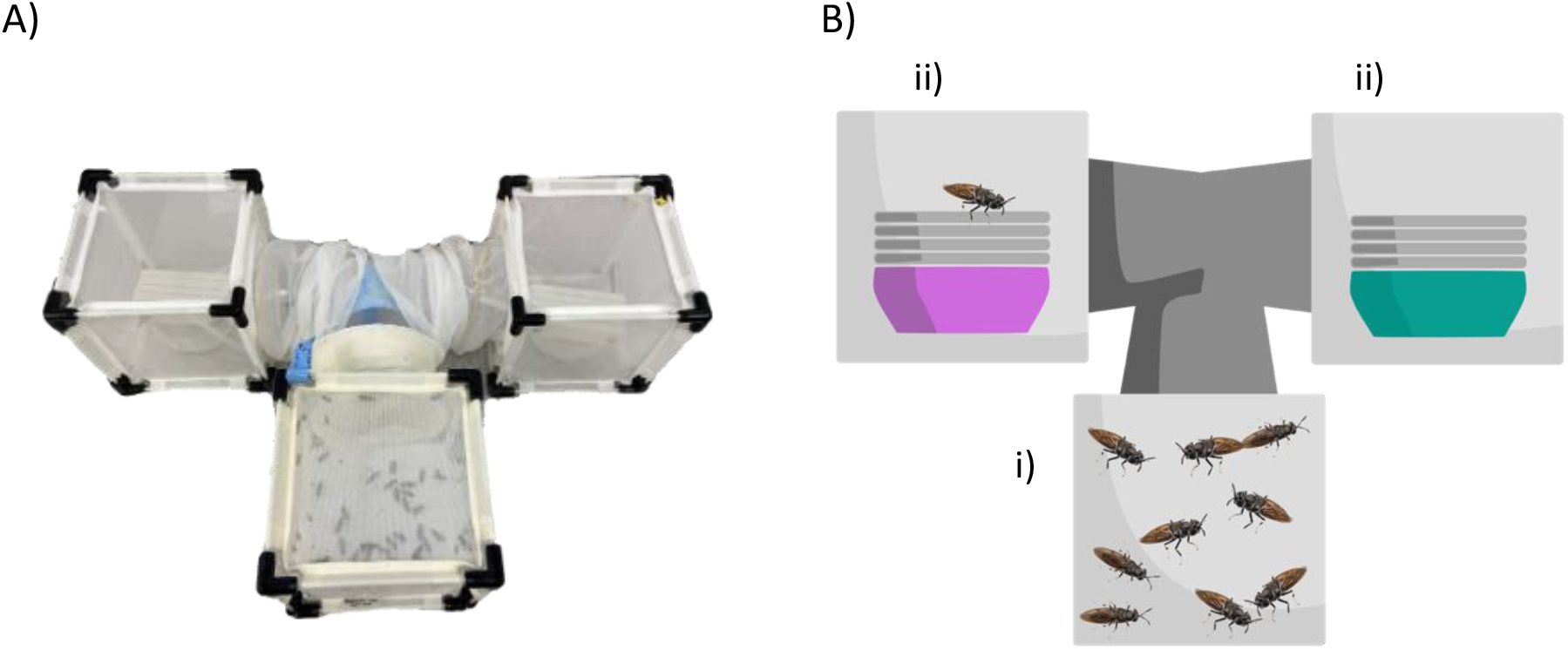
T-Maze Choice Assay Used in Behavioural Assays. A) Image of the T-Maze setup. B) Diagram of the T-Maze setup. Adult flies begin in the starting cage (i) and can travel between two cages (ii) containing different substrates, or a substrate and a control. Eggs are collected in a layered artificial acrylic/wooden egg trap, placed above a pot containing the substrate of interest.

To confirm candidate substrates as oviposition attractants, attractant substrates were placed in a T-maze assay against a solvent control (***Table 1***). The attractiveness of attractants was quantified by the mass of eggs laid per female per attractant:

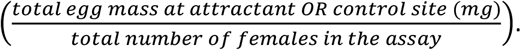

During behavioural assays using artificial compound mixtures, stock solutions of 100 mM of compounds were made by dissolving decanal, sulcatone, pentanoic acid, benzaldehyde, nonanal in hexane, and p-cresol, acetophenone, naphthalene, tetramethyl pyrazine, phenylethyl alcohol, 2-ethylhexanol, 2-methoxy-phenol, d-limonene, alpha-pinene, 1-octanol – in dichloromethane. Next, 10 ml of each of the 5 stock solutions were mixed together and 10 ml of DI water was added to the mixture. Dichloromethane and *n*-hexane, mixed in the same proportions as they occur in the compound mixture, and 10 ml DI water was added, served as solvent controls in these assays.

### Volatile collection

25 ml of the prepared putative attractants (***Table 1***) were placed in glass containers. The containers were enclosed in sealed oven bags (Bratschlauch, Toppits, Cofresco Frischhalteprodukte GmbH & Co. KG, Minden) and left for 30 minutes to allow volatiles to increase in concentration inside the bag.

Volatiles were collected with glass tubes (15 mm x 1.9 mm internal diameter) filled with Tenax (1.5 mg) and Carbotrap (1.5 mg) adsorbents. Both sides of these traps were closed with glass wool to keep the adsorbents in place (Otieno et al., 2023). Air was drawn from the oven bag through the trap with the help of a pump (DC12/16FK, Fürgut GmbH, Tannheim) for 15 minutes. Between four and eleven repeats of each attractant were collected. The traps were stored at 4 °C until transported to the University of Würzburg, Germany, where the samples were analysed.

### Gas chromatography/mass spectrometry analysis

To analyse the collected volatiles, we inserted each volatile trap into a glass-wool-packed thermodesorption tube and placed it in the thermodesorber unit (TDU; TD100-xr, Markes, Offenbach am Main, Germany). The thermodesorption tube was heated up to 260 °C for 10 min. The desorbed components were transferred to the cold trap (5 °C) to focus the analytes using N_2_ flow in splitless mode. The cold trap was rapidly heated up to 310 °C at a rate of 60 °C per second, held for 5 min, and connected to the gas chromatography/mass-spectrometry unit (GC–MS, Agilent 7890B GC and 5977 MS, Agilent Technologies, Palo Alto, CA, USA) via a heated transfer line (180 °C). The GC was equipped with an HP-5MS UI capillary column (0.25 mm ID × 30 m; film thickness 0.25 μm, J & W Scientific, Folsom, CA, USA). Helium was the carrier gas using 1.2 ml/min flow. The initial GC oven temperature was set at 40 °C for 1 min, then raised to 300 °C at 5 °C per min, where it was held for 3 min. The transfer line temperature between GC and MS was 300 °C. The mass spectrometer was operated in electron impact (EI) ionization mode, scanning m/z from 40 to 650, at 2.4 scans per second.

The relative abundance of the compounds in each chromatogram was calculated by integrating the relevant peaks. The ten most abundant compounds of each putative attractant were selected ignoring contaminants. Compounds eluting after 30 min were excluded from the analysis due to lack of volatility. Chemical compounds were identified using the NIST library and retention indices (Gold, 2019; Kováts, 1958; Kováts and Weisz, 1965). Following confirmation of compound identity, compounds were searched in Pherobase (El-Sayed, 2022) for known insect semiochemical activity. Finally, we calculated the relative composition of the selected compounds for all putative attractants.

### Electroantennography

Antennal responses to given odours were measured using an electroantennography (EAG) setup (Syntech, Kirchzarten, Germany). EAG was performed on sexually mature females (> 2 days post-emergence). Antennae were prepared by excision from BSF heads at the scape base, as well as removal of around 0.5 mm from the distal end of the flagellum with a sharp blade. Both sides of the antenna were inserted between two glass electrodes (ID 1.17 mm, Syntech), which were filled with Ringer solution (Ephrussi and Beadle, 1936). The antennal signal underwent a 10-fold pre-amplification, was converted into a digital signal using a DC amplifier interface (IDAC-2, Syntech), and was subsequently recorded using a GC-EAD software (GC-EAD 2014, version 1.2.5, Syntech). 1 μg doses of candidate compounds were tested. The compounds were diluted in dichloromethane (DCM) or hexane, as for the **Binary-choice behavioural assays** (see above). 10 μl of the dilutions (100 ng/μl) were loaded onto filter paper, acting as stimulant cartridges, and placed within glass Pasteur pipettes. Antennae were stimulated for 0.5 sec using a Stimulus Controller (CS-55, Syntech). The stimulation air (2 l/min) was directed into a consistently charcoal-filtered and humidified air stream (1 l/min). DCM or hexane served as the control stimuli. Each antenna was used to record 17 stimuli, including solvent control at the beginning and the end of the recording and repeated on 7 different antennae.

### Statistical analysis

Egg mass data and EAG data (**Figures 2, 4, 5**) are presented as median values, with error bars indicating interquartile ranges. Values were analysed with a Kruskal-Wallis test (with Dunn’s correction for multiple comparisons in Fig 4A and Fig 5).

**Figure 2.**
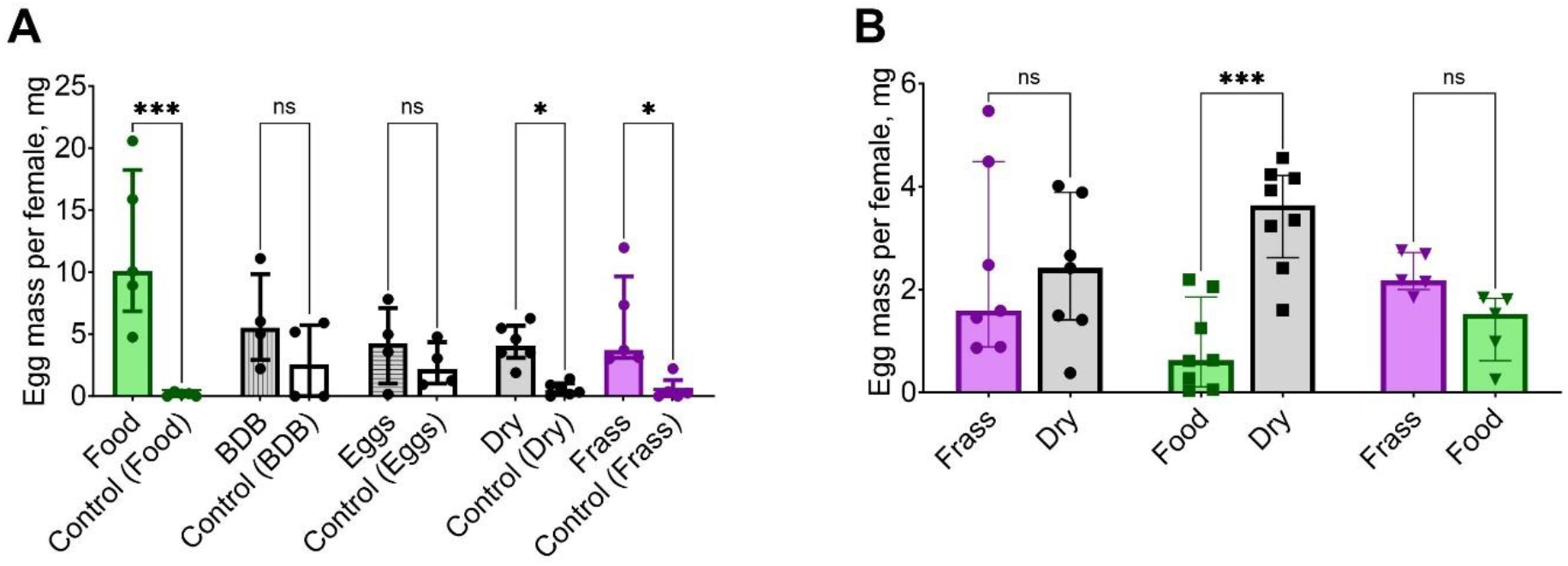
Responses to natural oviposition stimuli. A. Oviposition preference of stimuli vs control. Masses of eggs collected at the test substrate sides, and at their control sides. Food, Dry and Frass attracts significantly higher (Kruskal-Wallis test, ***p< 0.001, * p<0.05; ns – p>0.05) masses of eggs compared to their respective control sides. Food-Control, n=5; BDB-control, n=4; Eggs-Control, n=4; Dry-Control, n=6; Frass-Control, n=5. **B. Oviposition preference of stimuli vs each other**. In a separate set of experiments, the flies had to choose between two attractive oviposition substrates. Dry was significantly preferred over Food in this assay (Kruskal-Wallis test, *** p<0.001, ns – p>0.05). Frass-Dry, n=7; Food-Dry, n=8; Frass-Food, n=5.

**Figure 3.**
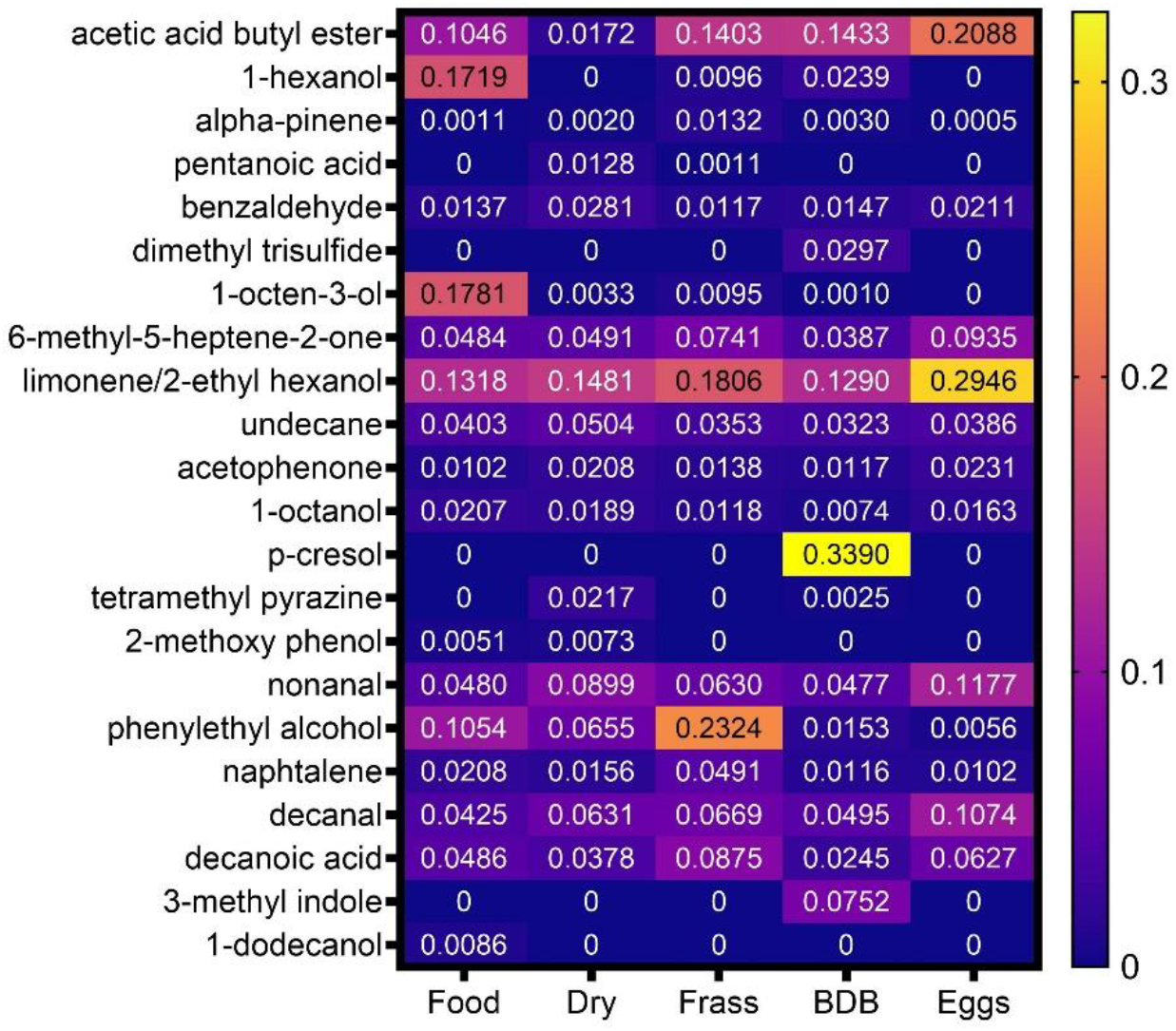
Chemical composition of volatiles in the headspaces of natural substrates. Average relative abundances of 23 most abundant volatiles, found in the headspaces of the 5 natural substrates. Food, n=4; Dry, n=7; Frass, n=6; BDB, n=11; Eggs, n=4.

**Figure 4.**
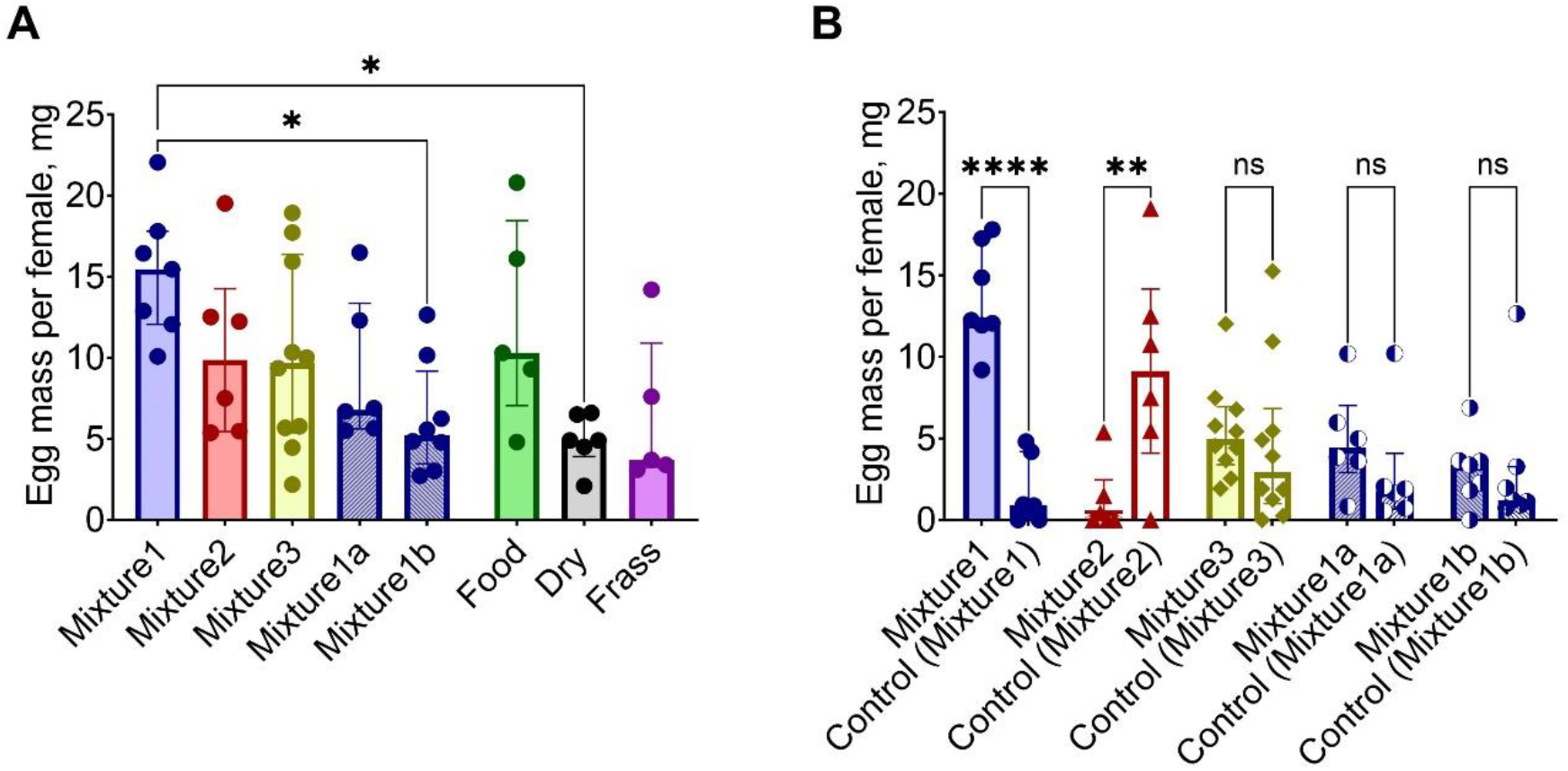
Behavioural responses to synthetic mixes. A. Total egg mass. The total mass of eggs collected in the choice assay (Food, Dry and Frass are re-plotted from Fig 2). Each data point represents the mass of eggs collected on the test side + the mass of eggs collected on the control side for the duration of the experiment, per female, in mg. Kruskal-Wallis test, * p<0.05. **B. Oviposition preference of stimuli vs control**. The same data as shown in **A**, but the masses of eggs collected at the test sides, and at their control sides, are shown separately. Mix1 attracted a significantly higher mass of eggs than its control (Kruskal-Wallis test, **** p<0.0001), and Mix2 attracted a significantly lower mass of eggs than its control (Kruskal-Wallis test, ** p=0.0029). Mix1 - Control, n=7; Mix2 - Control, n=6; Mix3 - Control, n=10; Mix1a - Control, n=6; Mix1b - Control, n=7.

**Figure 5.**
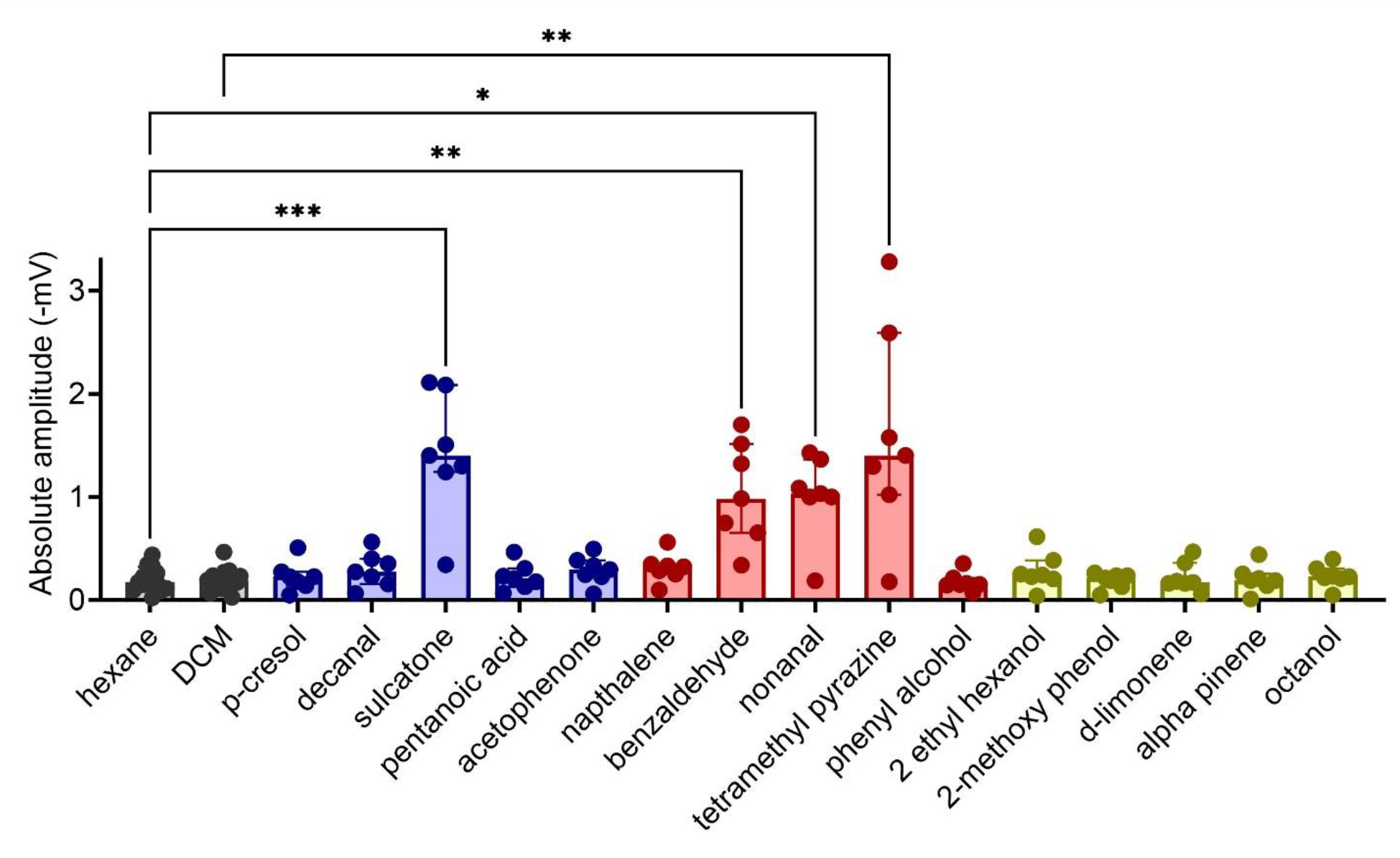
EAG responses of mature female BSF to compounds of interest. Compounds were tested at 1 µg dosage. Hexane and DCM were solvent controls. P-cresol, decanal, sulcatone, pentanoic acid and acetophenone (blue) were compounds of Mix 1; naphthalene, benzaldehyde, nonanal, tetramethyl pyrazine and phenyl alcohol (red) were compounds of Mix 2; 2-ethyl hexanol, 2-methoxy-phenol, d-limonene, alpha-pinene and octanol (yellow) were compounds of Mix 3. N=7 repeats for all test compounds, and n=14 for hexane and DCM controls. Bars depict median values, error bars depict first and third quartile ranges. Stars indicate a significant difference between a compound and its corresponding solvent control, Kruskal-Wallis test with multiple comparisons correction, * p<0.05, ** p<0.01, ***p<0.001.

Statistical analyses were conducted in the GraphPad Prism software v10.1.2. (324).

## Results

### Behavioural responses to volatiles of natural putative attractants

To determine the efficacy of various natural substrates in cueing oviposition, adult flies were provided with a binary choice assay, wherein gravid females chose between either an oviposition site containing a putative attractant, or a control (DI water for Food, BDB, Dry and Frass, and an empty container for Eggs). Putative attractants tested were fresh larval feed (Food), two types of larval waste and spent feed (Frass, Dry), anaerobically brewed larval and pupal remains (BDB) and conspecific eggs (Eggs) (see ***Table 1*** and **Materials and Methods** for details).

First, we asked whether each of the five substrates attracted more eggs per female than its corresponding DI water control. Food, Dry and Frass attracted significantly more eggs than their corresponding DI water control, and thus turned out to be effective oviposition attractants (***Fig 2A***, median M_Food_ = 10.1 mg, M_Food_control_ = 0.2 mg, p < 0.001; M_Dry_ = 4.1 mg, M_Dry_control_ = 0.5 mg, p = 0.03; M_Frass_ = 3.7 mg, M_Frass_control_ = 0.3 mg, p = 0.01). BDB and Eggs did not attract more eggs than their control sides, and thus did not act as effective oviposition attractants (median M_BDB_ = 5.5 mg, M_BDB_control_ = 2.6 mg, p = 0.13; M_Eggs_ = 4.3 mg, M_Eggs_control_ = 2.2, mg, p = 0.65), even though BDB and Eggs stimuli induced the flies to lay relatively high amounts of eggs in total.

This assay demonstrated that Dry, Food and Frass act as oviposition attractants. Thus, we compared these three attractants directly in binary choice assays to investigate their relative attractiveness.

When tested in a t-maze choice assay (***Figure 2B***) females strongly preferred Dry over Food (M_Dry_ = 3.6 mg, M_Food_ = 0.6 mg, p < 0.001). However, preference for neither of the two tested attractants was significant in the tests with Frass vs Dry and Frass vs Food stimuli (M_Frass_ = 3.6 mg, M_Dry_ = 0.6 mg, p = 0.96; M_Frass_ = 3.6 mg, M_Food_ = 0.6 mg, p = 0.11).

Next, we investigated the chemical composition of volatiles, emitted by each of the five substrates.

### Identification of compounds in the headspace of attractants

As oviposition is triggered by an olfactory cue, the variation in substrate attractiveness must stem from the unique volatile bouquets emitted from different attractants. We have identified the 23 most abundant compounds in the headspace of the 5 natural substrates (***Figure 3***) that came from various chemical classes: alcohols, aldehydes, esters, fatty acids, aromatic compounds, terpenes etc. We were not able to quantify the abundance of limonene separately from 2-ethyl hexanol, as both of these compounds occurred in the headspaces of all substrates tested, and their respective peaks overlapped in the chromatograms. 13 of the 23 compounds occurred in the headspace of all natural substrates, and only 5 were unique for their substrates. The most attractive substrate, Dry, and another attractive substrate, Frass, had no unique compounds in their headspaces. The third attractive substrate, Food, had a unique compound 1-dodecanol, but at a relatively low abundance of 0.86%. 2-methoxy phenol and pentanoic acid occurred in two out of three attractive substrates and have not occurred in the headspace of non-attractants. Tetramethyl pyrazine occurred in both the strongest attractant, Dry (2.17%), and a non-attractant BDB (0.25%). A non-attractant BDB had two unique compounds, a highly abundant p-cresol (34%) and dimethyl trisulphide (3%), while Eggs had no unique compounds. Since no compounds clearly accounted for the difference between attractants (Food, Dry, Frass) and non-attractants (Eggs, BDB), we have reasoned that the differential attractiveness of natural substrates is due to the specific combinations of the common and unique compounds in their headspace, rather than the presence of specific unique compounds.

To explore the attractiveness of naturally occurring compounds, we decided to test them as artificial mixtures. We selected 15 semiochemically active compounds that occurred in Food, Dry and Frass attractants (***Figure 3***) at relatively high (decanal, nonanal, phenylethyl alcohol, limonene/2-ethyl hexanol, sulcatone (6-methyl-5-heptene-2-one)) and low (naphthalene, 1-octanol, acetophenone, benzaldehyde, alpha-pinene) abundances, as well as compounds that occurred in only some of the substrates (2-methoxyphenol, tetramethyl pyrazine, p-cresol, pentanoic acid).

### Behavioural responses to artificial mixes

To test the oviposition attractiveness of 15 selected compounds, we created 3 mixtures, each containing 5 of the 15 compounds:

**Mixture 1:** p-cresol, decanal, sulcatone, pentanoic acid, acetophenone

**Mixture 2:** naphthalene, benzaldehyde, nonanal, tetramethyl pyrazine, phenylethyl alcohol

**Mixture 3:** 2-ethylhexanol, 2-methoxyphenol, d-limonene, α-pinene, octanol

The mixtures were tested in the binary choice assay (see **Materials and Methods** for details).

First, we investigated the total mass of eggs laid per female in the three assays. Mixture 1 assays elicited the largest overall egg mass of the three mixes tested (median: 15.5 mg, 1^st^ quartile: 12.1 mg, 3^rd^ quartile: 17.8 mg). Mixture 2 (median: 9.9 mg, 1^st^ quartile: 5.5 mg, 3^rd^ quartile: 14.2 mg) and Mixture 3 (median: 9.7 mg, 1^st^ quartile: 5.4 mg, 3^rd^ quartile: 16.4 mg) elicited similar total masses of eggs per female. The total egg mass was not significantly different between the three assays.

Interestingly, the total mass of eggs per female laid in the assays with Mixture 1 was significantly higher than that in assays with the most attractive natural stimulus, Dry (p = 0.02, ***Figure 4A***).

Second, we investigated whether the artificial mixes act as oviposition attractants. Mixture 1 was significantly more attractive than its control, with most of eggs within the assay laid in the oviposition site containing the mix (median M_Mix1_ = 12.3 mg, M_Mix1_control_ = 0.9 mg, p < 0.001, ***Figure 4B***). To test if one of the compounds of Mixture 1 is sufficient to cue oviposition, Mixture 1 was split into two mixtures:

**Mixture 1A**: decanal, p-cresol

**Mixture 1B:** sulcatone, pentanoic acid, acetophenone.

When tested, neither Mixture 1A nor 1B proved significantly attractive (median M_Mix1A_ = 4.4 mg, M_Mix1A_control_ = 1.8 mg, p = 0.25; median M_Mix1B_ = 3.4 mg, M_Mix1B_control_ = 1.2 mg, p = 0.79, ***Figure 4B***). The lack of ovipositing attractiveness when Mixture 1 was split indicates that not one compound, but a mixture of at least two compounds is necessary to cue oviposition. Additionally, the total mass of eggs laid per female during assays utilizing either Mixture 1A (median: 6.8 mg, 1^st^ quartile: 5.6 mg, 3^rd^ quartile: 13.4 mg) or Mixture 1B (median: 5.2 mg, 1^st^ quartile: 3.5 mg, 3^rd^ quartile: 9.2 mg) was much lower than mass of eggs in Mixture 1 assays, with Mixture 1B assays producing significantly lower mass of eggs than Mix 1 (p = 0.047, ***Figure 4A***).

Mixture 2 turned out to be a strong oviposition repellent, with most of the eggs laid within the assay deposited at the solvent control ovipositing site (median M_Mix2_ = 0.2 mg, M_Mix2_control_ = 9.1 mg, p = 0.003).

Mixture 3 has not acted as an oviposition attractant or repellent. There was no significant difference between the mass of eggs laid per female at the Mix 3 site and its control (median M_Mix3_ = 5.0 mg, M_Mix3_control_ = 2.9 mg, p = 0.28) (***Figure 4B***).

To investigate the basis of the observed behavioural responses to artificial mixtures (***Figure 4***), we next asked whether the adult female flies could smell the 15 compounds that comprised the mixtures.

### Sensory detection of volatile compounds

To test whether the flies can detect our selected 15 compounds, we performed electroantennography on female BSF, presenting compounds at 1μg dosage.

Female antennae responded at a range of 0 – 3.3 mV to compounds of interest (***Figure 5***). Responses to sulcatone, benzaldehyde, nonanal and tetramethyl pyrazine were significantly different from corresponding solvent controls. Interestingly, only one of these compounds, sulcatone, was included in attractive Mixture 1, while the other 3 compounds were included in Mixture 2, and none of the compounds that elicit strong antennal responses were included in Mixture 3.

In summary, we identified an effective synthetic oviposition attractant, Mixture 1, and an oviposition repellent, Mixture 2, and established the responses of the BSF peripheral olfactory system to its constitutive compounds. Interestingly, our results also indicate that the oviposition attractiveness of natural or synthetic mixtures is not linked to the total egg mass laid by the flies when exposed to these mixtures.

## Discussion

This study achieved its original aim to formulate a synthetic oviposition attractant for Black Soldier Flies. Below we discuss additional lines of interest that arose during this study and will require further investigations.

### Olfactory cues may be oviposition attractants or repellents

None of the five natural substrates that we investigated (*Figure 2A*) acted as oviposition repellent. However, a synthetic mix of pure compounds (Mixture 2, *Figure 4B*) deterred oviposition. This result implies that natural volatile mixtures may also repel the oviposition of BSF. Indeed, the bacterial content of oviposition substrates may serve both as an oviposition attractant and repellent, depending on the concentration and species of bacteria (Zheng et al., 2013). It is possible that this response was also mediated, at least in part, by the BSF olfactory system via the perception of bacteria-derived volatiles. Olfactory oviposition attractants and repellents have been identified in multiple other insect species. For example, in *Drosophila melanogaster*, volatiles from citrus fruit serve as olfactory oviposition attractants (Dweck et al., 2013), and geosmin – as oviposition repellent (Stensmyr et al., 2012). Interestingly, acetic acid serves as a gustatory oviposition attractant, and, simultaneously, as an olfactory oviposition repellent, in *D. melanogaster* (Joseph et al., 2009). It is also possible that BSF oviposition may be controlled via multiple chemical compounds that are detected by gustatory and olfactory systems.

### What makes an olfactory stimulus an attractant or a repellent?

Olfactory coding and other neuroscience phenomena have not been investigated in BSF in any detail so far. However, extensive research conducted on *D*.*melanogaster* and other organisms points towards multiple mechanisms valence may be encoded in the peripheral olfactory system. First, a “labelled line” mechanism elicits a behavioural response whenever a certain single type of olfactory sensory neurons is activated by their chemical ligand (Chin et al., 2018; Kurtovic et al., 2007; Stensmyr et al., 2012). Usually, labelled lines respond to highly ecologically important compounds, such as pheromones or highly toxic substances. Second, repellency may be also elicited by high concentration of many chemicals, presumably via general over-activation of the olfactory system, which may be seen as a variation of the labelled line mechanism (Stensmyr et al., 2003; Yoshida et al., 2012). Third, activation of certain specific groups of sensory neurons may also be interpreted in the brain as an attractive or a repellent stimulus, whereas activation of a subgroup of these neurons will not lead to the same behavioural response. This last mechanism is known as combinatorial coding (Haverkamp et al., 2018).

Our electrophysiological experiment indicated that BSF can detect only one compound, sulcatone, of attractive Mixture 1 (*Figure 5*). However, the oviposition assays with Mixtures 1, 1A and 1B show that the attractiveness of Mixture 1 cannot be explained by one compound only, otherwise Mixture 1B which contains sulcatone would have also been attractive. Sulcatone thus does not activate a “labelled line” response in BSF, and other volatiles in combination with sulcatone are needed to produce an oviposition attractant. Why was there then no electrophysiological response to other compounds of Mixture 1? The EAG responses reliably convey the simultaneous activation of many sensory neurons. If only a handful of neurons are active, the EAG signal is likely to be weak and undetectable from noise. It is thus likely that other behaviourally active components of Mixture 1 activate only a few olfactory sensory neurons, while sulcatone activates many.

Mixture 2 is an oviposition repellent, and 3 components of it elicited EAG responses. Each of the three mechanisms described above is equally likely to account for repellency in this case. Further experiments are necessary to establish the exact mechanism of repellency of Mixture 2.

ORCO-dependent olfactory receptors expressed in BSF are known (Xu et al., 2020), and the first genetically modified BSF have been generated recently (Gunther et al., 2024). It now becomes feasible to conduct detailed genetic investigations and manipulations of the BSF olfactory system to establish receptor-ligand pairs and neuron-behaviour connections. Generation of new BSF lines with genetically altered behavioural preferences is also now feasible and may facilitate their mass rearing for industrial purposes.

### Oviposition repellent does not suppress egg laying in BSF

We observed that Mixture 2, while being a strong oviposition repellent, doesn’t suppress egg laying in BSF, but rather spatially redirects it (Figure 4). This is an important observation, as Mixture 2 may thus also be used in industrial settings, perhaps in combination with Mixture 1, without detriment to overall egg production. While the exact mechanism of olfactory enhancement or suppression of oviposition is not clear, evidence from other organisms indicated that oviposition repellents often also suppress overall egg production. For example, CO_2_ is a behavioural repellent and suppresses egg production in *C. elegans* (Fenk and de Bono, 2015). Rotten fruit, when compared to ripe fruit, both repel oviposition and suppress egg laying in *D. suzukii* (Karageorgi et al., 2017), while ripe cherry, strawberry and banana are both oviposition attractants when compared to other ripe fruit, and induce flies to lay more eggs (Cai et al., 2019). Similarly, phenol is both an oviposition repellent and suppresses egg production in *D. melanogaster* (Mansourian et al., 2016). Further studies of how olfactory stimuli direct oviposition and control egg production in BSF are required to understand why BSF is different in this respect from other organisms.

### What affects the olfactory preferences of BSF?

In this study, we evaluated the innate oviposition preference of BSF. However, based on previous reports, this preference is not fully genetically hardwired and may be affected by the food that flies received during their larval stage. Natal habitat preference has been previously reported for both insects and mammals (Akhtar and Isman, 2003; Anderson et al., 1995; Merrick and Koprowski, 2016; Moreau et al., 2008). Our preliminary data also indicated that supplementing BSF larval chick-crumb feed with bananas increases preference for banana volatiles in oviposition substrate (*NKTT and OR, unpublished*). The extent of these effects requires further investigation.

### Future work

This project was driven by an industrial need to have an easy-to-make and easy-to-distribute reproducible synthetic oviposition attractant, none of which were previously reported. The attractant and repellent we formulated have so far been tested in small-scale research laboratory settings. Immediate future steps for industrial purposes now include: 1) large-scale industrial tests; 2) test of a range of concentrations to identify optimal ones; 3) test of mixtures, derived from Mixture 1 and 2, but with a reduced number of compounds, to optimize mixture composition and cost; 4) tests of other suitable solvents; 5) engineering of easy to distribute formulations, perhaps in the form of gels or pellets.

Our results also highlighted the paucity of knowledge about BSF sensory systems. While an industrially important insect, BSF is an ideal new model system for studies of brain anatomy and function, due to its large size, ease of rearing and maintenance, and the possibility of genetic modifications. Fieldwork or lab work under outdoor conditions may be conducted on BSF in the tropics and sub-tropics, and our preliminary observations indicate that there is much to discover from these future studies.

## Acknowledgements

We thank Elaine Fitches for encouraging us to develop this project and allowing us to use her insectary, and Sam Grainger, Thomas Farrugia and the entire BetaBugs team for supplying us with flies and training us in BSF maintenance. We thank Vincent Croset and Sonia Sen for helpful comments on the manuscript.

## Author contributions

OR and TS conceived the project; NKTT, ZK, TS and OR designed experiments; NKTT performed all experiments; NKTT and ZK analysed data; all authors co-wrote the manuscript.

## Data availability

All original data not included in this manuscript are available from corresponding authors upon request.

